# Insights into Early Word Comprehension: Tracking the Neural Representations of Word Semantics in Infants

**DOI:** 10.1101/2021.10.28.466334

**Authors:** Rohan Saha, Jennifer Campbell, Janet F. Werker, Alona Fyshe

## Abstract

Infants develop foundational language skills and can understand simple words well before their first birthday. This developmental milestone has been studied primarily using looking time paradigms and Event-Related Potential (ERP) techniques, which provide evidence of word comprehension in the infant brain. While prior research validates the presence of semantic representations of words (word meaning) in infants, it does not tell us about the mental processes involved in retrieving these semantic representations or the content of the representations. To this end, we use a machine learning approach to predict the semantic representations of words using Electroencephalograms of infant brain activity. We explore semantic representations in two groups of infants (9-month-old and 12-month-old) and find significantly above chance decodability of overall word semantics, word animacy, and word phonetics. We compare decodability between the age groups and find significant differences later in the analysis window (around 700-900 ms after word onset), with higher decoding performance in 9-month-olds. Through our analysis, we also observe strong neural representations of word phonetics in the brain data for both age groups, some possibly correlated to word decoding accuracy and others not. Finally, we use the Temporal Generalization Method to show that neural representation of word semantics generalizes across the two infant age groups. Our results on word semantics, phonetics, and animacy provide insights into the evolution of neural representation of single word meaning in infants.

## 1 Introduction

Studies using looking time paradigms and Event-Related Potentials (ERP) techniques have provided compelling evidence indicating that infants understand the meaning of multiple words by six months of age, as demonstrated by their ability to associate known words with corresponding objects [Bergelson and Swingley, 2012, Tincoff and Jusczyk, 2012]. While these investigations have contributed significantly to our understanding of word semantic comprehension in infants, there exists a notable gap in the literature concerning the characterization of the neural representations themselves.

Our present study aims to address this gap by delving into the dynamics of neural representations in infants concerning various aspects of word meaning. Specifically, we investigate the differences in these neural representations between 9-month-old and 12-month-old age groups, focusing on single word meaning. Although research on the neural representations of word semantics in adults is extensive [Fyshe et al., 2019, Honari-Jahromi et al., 2021], the same is less true for infants. Our attention is directed towards unraveling the transient nature of these representations and discerning specific dissimilarities and similarities across the two infant age groups.

Infants’ early recognition of the correspondence between highly familiar words and objects is well-established [Tincoff and Jusczyk, 1999, Parise and Csibra, 2012, Bergelson and Swingley, 2012, 2015]. As infants encounter new words and objects, their vocabulary expands [Bergelson and Aslin, 2017]. While prior research has predominantly employed looking time paradigms to demonstrate the emerging ability of word understanding in infants [Bergelson and Swingley, 2012, Campbell and Hall, 2022, Arias-Trejo and Plunkett, 2010], the nature of the associated neural representations remains less explored.

Looking time paradigms conventionally leverage the N400 ERP component to investigate the mismatch between spoken or written words and their referents [Kutas and Federmeier, 2000]. The N400, a negative signal peaking around 400ms after stimulus onset, is indicative of semantic processing and reliably emerges in infants by one year of age [Friedrich and Friederici, 2010]. Notably, the N400 component has been employed to demonstrate that 9-month-old infants can detect a mismatch between an object and the preceding word label [Parise and Csibra, 2012]. In our study, we further probe the neural representations around 400ms after stimulus onset and throughout the duration of the trial, exploring the differences between 9-month-old and 12-month-old infants. To unravel the nature of these representations, we presented audio-visual stimuli to infants and collected their EEG data during stimulus presentation. Using a machine learning-based decoding model, we predicted word vector representations, to gain insights into the evolving dynamics of neural representations over time.

Machine learning has been widely applied to various neuroimaging modalities to study the neural processing of language in adults [Pereira et al., 2009]. These include applying encoding models to fMRI data to predict voxel activations of fMRI responses, to map various semantic domains onto brain areas [Huth et al., 2016a], applying decoding models on MEG data to explore the temporal changes of neural representations of nouns [Sudre et al., 2012], and also, to explore the semantic decodability of phrases [Fyshe et al., 2019]. Moreover, other modalities such as EEG data have also been successfully used with linear models to understand the neural processes involved in language learning in adults [Foster et al., 2021]. These studies suggest that machine learning can be used to study the dynamics of the neural representations of infants over time and give us insights into the onset of signals pertaining to single word semantics.

Similarly, previous works on infants have used various neuroimaging modalities, such as EEG and functional near-infrared spectroscopy (fNIRS). EEG data from infants are especially valuable because it helps us understand the temporal changes in the neural representations during the stimuli presentation. On the other hand, fNIRS is a technique that measures changes in the cortical blood oxygenation can provide with good spatial resolution, but lacks good temporal resolution. Such modalities can be analyzed using Multivariate Pattern Analysis (MVPA) techniques to study the activations involved in semantic processing of information.

Methods such as Support Vector Machine (SVM) and Convolutional Neural Network (CNN) have been successfully used to study aspects of language in infants, such as the classification of rhythmic stimuli from EEG data in 8-week-old infants [Gibbon et al., 2021]. And fNIRS data has also been successfully used to classify a set of stimuli into audio or visual categories [Emberson et al., 2017]. The usefulness of EEG data is also apparent in studies where the onset of specific effects is crucial. For example, EEG has been used to explore the developmental trajectory of visual category representations in infants younger than 1-year-old, and how it compares to those of adults [Xie et al., 2022]. The authors showed that the temporal dynamics of visual category representations change from slowly emerging simple representations to rapidly evolving complex features in adults. EEG has also been used to study onset of certain neural effects involved in information processing. Another study used Support Vector Machine (SVM) model to correctly decode and classify images of animals and body parts from EEG data [Bayet et al., 2020], and also showed that this decodability was observed within 500ms of stimulus onset. Such works motivate the investigation of the differences of the neural representations of young infants of 9- and 12-month-old, and the onset of the signals related to semantic processing of single word.

To summarize, similar to previous decoding studies on adult neuroimaging data [Huth et al., 2016b, Fyshe et al., 2019, Honari-Jahromi et al., 2021], we use a decoding model on EEG data collected from 9-month-old and 12-month-old infants to predict word vector representations of single words.

The following are the contributions of our study.

- Our analysis shows high decoding accuracy of words immediately after word onset.
- Non-semantic word stimuli properties such as phonetics can be decoded from infants of both age groups.
- The neural responses of 9-month-old and 12-month-old infants show similar representations of word semantics across time windows.
- The neural representations of word semantics in both groups of infants show signs of generalizability across different time points.

## 2 Materials and Methods

### 2.1 Participants

A total of 46 infants participated in the study. Of this initial sample, twenty-one 9-month-old (10 female, 11 male) infants were included in the final analysis. Seven additional 9-month-olds were tested but excluded from the final sample due to fussiness (*n*=5), poor electroencephalogram (EEG) impedance (*n*=1), and excessive body movements (*n*=1). Fourteen 12-month-old (7 female, 7 male) infants were included in the final sample. An additional four 12-month-old infants were tested but were excluded due to language criteria (*n*=1, the majority of the language spoken at home was not English) and fussiness (*n*=3). All infant participants were native English learners, had no cognitive impairments, and were born full-term.

### 2.2 Stimuli

The stimuli presented to the infants consisted of two components, audio stimuli, and visual stimuli. The audio stimuli set consisted of sixteen total words equally divided into animate and inanimate words. The stimuli words were selected based on the reported age of acquisition from the MacArthur–Bates Communicative Development Inventory (CDI) [Fenson, 1993] and numerous other experimental studies [Bergelson and Swingley, 2012, Tincoff and Jusczyk, 1999, 2012, Bergelson and Swingley, 2015, 2018]. A grouping of the stimuli words based on animacy is presented in Table 1.

**Table 1:**
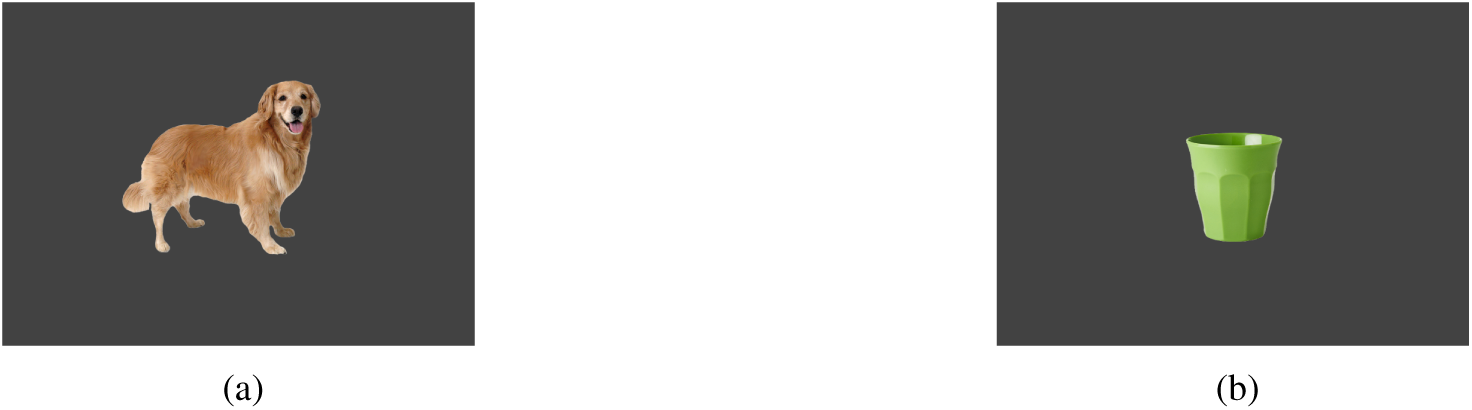
Stimuli words used in the study.

The visual stimuli consisted of sixteen images for each stimulus word. Each image was publicly available on Google Images and had black backgrounds, and represented a photographic depiction of the corresponding word label and not toys. For example, the image for *dog* showed a real-life dog and not a toy dog. An example of an animate and an inanimate image is provided in Fig 1.

**Figure 1:**
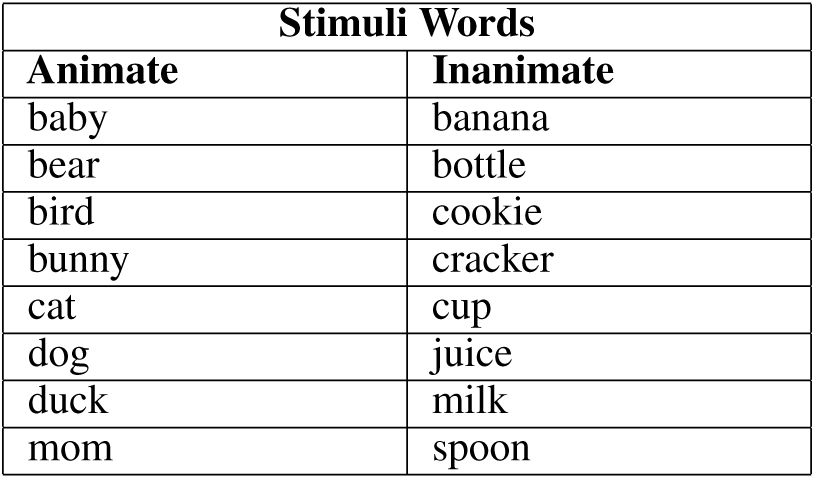
An example of dog (a) as an animate (a) and a cup (b) as an inanimate image presented to the participants after the word stimuli.

### 2.3 Word vectors

We represented the word stimuli using word vector representations obtained from the Word2Vec model [Mikolov et al., 2013], which is pretrained on the Google News Dataset (an internal Google dataset containing news articles and comprising about a hundred billion tokens^2^). These word vectors represent the semantic properties of the stimuli. Each word vector has 300 dimensions. For our study, we used the vectors acquired from the skip-gram algorithm, a neural network trained to predict the context for a given word. We used vectors from Word2Vec because several existing studies show that Word2Vec representations of stimuli can be decoded from various brain-imaging modalities in adults [Ruan et al., 2016, Kivisaari et al., 2019, Foster et al., 2021, Honari-Jahromi et al., 2021].

### 2.4 Data collection and Preprocessing

We recorded the infant participants’ neural data using Electroencephalography (EEG). To record the EEG, we used a Geodesic 64 channel EEG cap which had a data capturing frequency of 1000 hertz. We removed channels 61-64 during preprocessing since these channels corresponded to cheek channels and were not present on all caps. We used Eprime software with Chronos hardware [Schneider et al., 2002] to carry out the experiment, and EGI Netstation 5.4.2 software to capture data.

Before each trial, a computer displayed a dynamic video as an attention getter on a screen placed in front of the participants who were sitting on their parent’s lap. The video displayed a small rotating asterisk that changed colors and was used to ensure the participants’ attention based on the criteria that they were either looking at the rotating asterisk or the experimenter. After ensuring a participant’s attention, the Eprime software presented a stimulus word through the experimenter’s (a female native English speaker) headphones (to ensure that the infants and parents could not hear the stimulus word), who then spoke the word into a microphone signalling the word stimulus onset. To control for potential differences in pronunciation of the stimuli words, only one person (the experimenter) spoke all the stimuli words in the microphone for all participants. Following the word stimulus onset, the rotating asterisk was shown for another 1100ms. The Eprime software then displayed the image stimulus on the screen (Image Onset) that lasted for 1200ms (Image Offset). After the image offset, a blank screen was shown for 1000ms. We show the timing diagram in Fig 2.

**Figure 2:**
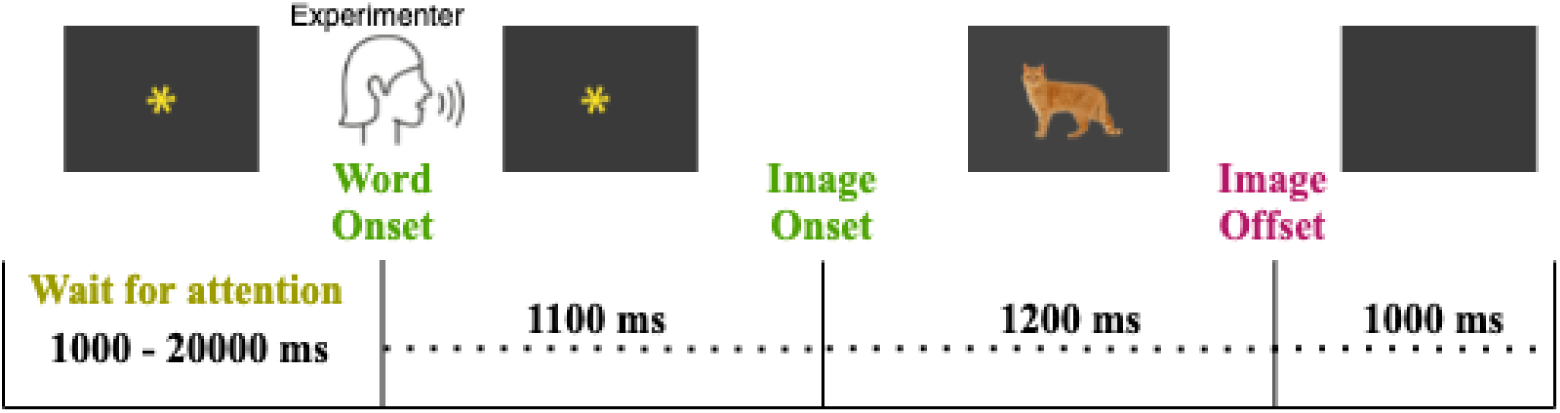
Timing diagram showing the stimulus presentation design. The word stimulus was in audio format spoken, and the image was shown on a screen. Before each trial, a dynamic video was shown with a rotating asterisk that changed colors. After word onset, the rotating asterisk was shown for at least 1100ms. Once the infant fixated on the screen, an image appeared on the screen lasting 1200ms. After the image offset, a blank screen followed that lasted 1000ms.

For the software to register the onset of the spoken word stimulus, a threshold level of 65dB was used. Afterward, we manually corrected any inconsistencies in the word onset. On average, the time between image offset and the next word stimulus varied from 2100ms to 2500ms because participants’ attention was required, and the reaction time of the speaker was accounted for to the next word stimulus provided by the Eprime software. During the image presentation, for half of the trials, the image matched the word label (for example, an image of *dog* was shown for the preceding word label *dog*). For the other half of the trials, the image did not match the preceding word label (for example, an image of a *cup* was shown for the preceding word label *banana*). Half of the mismatched images matched the animacy category of the previous word, and the remaining half mismatched the animacy category of the word. In all the analyses discussed in this paper, we only used the EEG data recorded during the word stimuli presentation as we were interested in exploring the change in the neural patterns of semantics over time. We do not use the EEG data recorded during the image presentation in any of our analyses.

For the analyses, we discarded trials where the participant was not paying attention (was not looking at the screen or the experimenter). We also removed any trials that contained movement artifacts, which were found by visual inspection of the EEG signal and the video feed of the infant’s face. After removing the bad trials, there were around 46 trials per participant in the 9-month-old group, and around 47 trials per participant in the 12-month-old group (see Fig S2 in the Supplementary Materials for more information). In the end, there were 974 trials of EEG data for the 9-month-old babies and a total of 665 trials of EEG data for the 12-month-old babies. The data were sampled at a rate of 1000Hz with no subsequent downsampling. Using EEGLAB v2022.0 [Delorme and Makeig, 2004] in MATLAB (version R2022a), we filtered the data using a high pass filter of 0.1Hz and a low pass filter of 50Hz, re-referenced the data using the average reference of all data channels, and extracted epochs from −200 to 1000ms at the onset of each word utterance (0ms). Finally, we baseline corrected the data from −200ms to 0ms.

### 2.5 Prediction Model

To predict the vector representations of the word stimuli (*Y*) from the EEG data (*X*), we used a decoding framework by using a ridge regression model from scikit-learn [Pedregosa et al., 2011]. The decoding framework has been successfully used in previous work to decode word vector representations from brain-imaging data [Sudre et al., 2012, Honari-Jahromi et al., 2021, Foster et al., 2021].

For all the *N* trials recorded for each age group, we chunk the EEG data of each trial into 100ms windows with a 10ms step size. Each chunked trial of size 100ms has data from all the 60 sensors. We then reshape this trial into a vector of size *p* = 60 *×* 100 = 6000 (sensor x time points). We follow this vectoring procedure on the data from all the windows of the trial. Repeating this procedure for all the *N* trials results in a matrix *X* of size *N × p* (see Fig 3).

**Figure 3:**
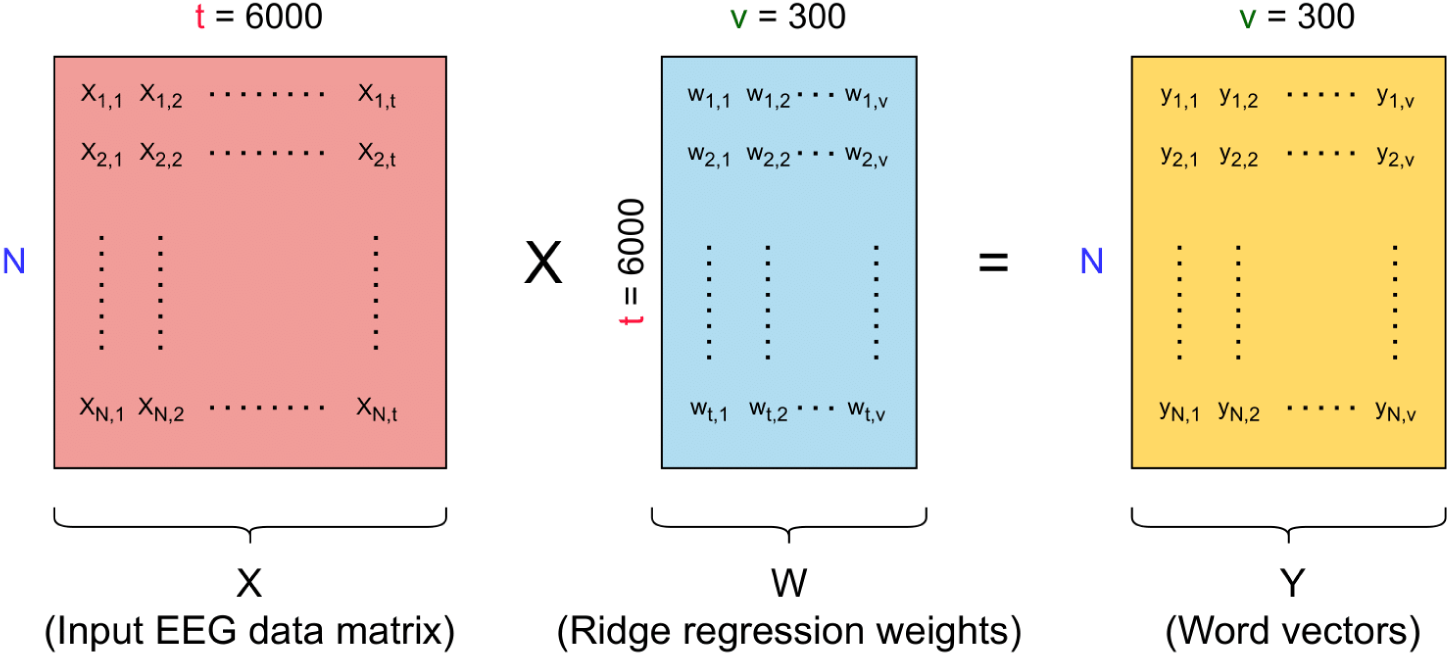
The ridge regression model takes the EEG data matrix as input and predicts the Word2Vec vectors. The input EEG data matrix *X* is of size *N × t*, where *N* is the number of trials and *t* is the number of EEG time points (across p=60 sensors and 100ms window size). The ridge regression model learns a weight matrix *W* of size *t × v*, where *v* is the number dimensions of Word2Vec embeddings. Finally, the ridge regression model predicts the Word2Vec vectors *Y* of size *N × v*.

We then divided the data matrix *X* into a training set *X_train_* with the corresponding vector representations *Y_train_* and a testing set *X_test_* with the corresponding vector representations *Y_test_*. We trained the regression model on *X_train_* to predict *Y_train_*, and evaluated its performance on *X_test_* and *Y_test_*. We used an 80-20 split, where 80% of the trials from the matrix *X* were used for the training set and 20% were used for the test set. There was no overlap between the trials of the two sets to avoid overfitting. We repeated this process fifty times (Monte-Carlo iterations).

To increase the signal-to-noise ratio, we averaged all the trials for a stimulus word in the test set *X_test_*. For example, if there are approximately 12 trials for the word ‘bottle’ in the test set, we averaged these 12 trials to obtain one ‘averaged trial’ for that word. These trials can come from multiple participants. For instance, in the 9-month-old group, the maximum percentage of data for the word ‘bottle’ that can come from one participant is 14.5% of the total number of trials for that stimulus word (see Fig S1 in the Supplementary Materials for the maximum percentage contribution of each word from a participant in both age groups). Thus, the trials for the stimulus words are distributed across multiple participants. To obtain the final test set, we repeated the averaging process for all the stimulus words, which resulted in 16 total trials in *X_test_*.

The Ridge Regression model is a mapping *W* that transforms the EEG data *X* to a prediction for the word vector representations *Y*. To learn this transformation, we trained the weights (**ŵ**) of the Ridge regression model by minimizing the loss function as follows.

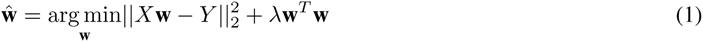

The term *λ***w***^T^* **w** is a ridge or the L2 regularize, and the hyperparameter *λ* controls the regularization strength. We used nested K-fold cross-validation (k=5) procedure with negative mean squared error to optimize the hyperparameter *λ* for the ridge regression model, and we used scikit-learn’s GridSearchCV^3^ to achieve this. In Fig 3, we show the weights of the Ridge Regression model being multiplied with the input EEG data (*X*) to predict the word vectors (*Y*).

### 2.6 2 versus 2 Test

To evaluate the performance of the model, we used the 2 vs 2 test procedure to measure the similarity of predicted word vectors to the ground truth word vectors. The test works by calculating the sum of the distances of the matching pairs of word vectors (*d*(*y_i_, ŷ_i_*) + *d*(*y_j_, ŷ_j_*)) and comparing that with the sum of the distances between the non-matching pairs of the word vectors (*d*(*y_i_, ŷ_j_*) + *d*(*y_j_, ŷ_i_*)). We used cosine distance as the distance metric.

The 2 vs 2 test passes if:

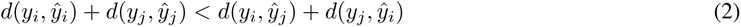

The 2 vs 2 test is depicted in Fig 4. The chance accuracy of the 2 vs 2 test is 50% because there are two possible assignments of the predicted vectors to the ground truth vectors (red and green lines shown in Fig 4), and the assignment of each predicted vector to each ground truth vector is equally likely. We consider all possible pairs. The total number of pairs evaluated by the 2 vs 2 test is 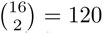, as each trial in the test set is the average across all the trials for a stimulus word in the test set (see Section 2.5).

**Figure 4:**
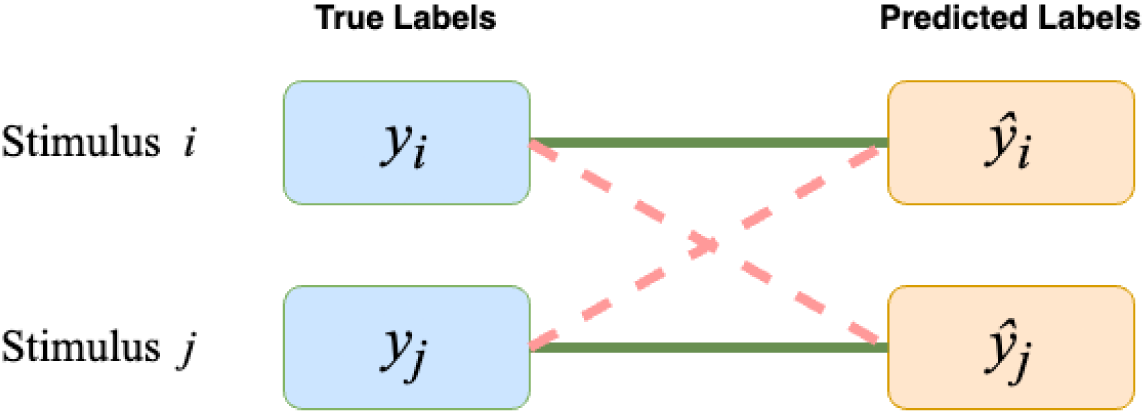
2 vs 2 test showing pairwise comparisons of true (*y_i_, y_j_*) and predicted labels (*ŷ_i_, ŷ_j_*). The 2 vs 2 test passes if the sum of the cosine distance between matching word vector pairs (solid green lines) is less than the sum of the distance between the non-matching word vector pairs (red dashed lines).

### 2.7 Testing for Above Chance Accuracy

To compute statistical significance, we used the permutation test [Ojala and Garriga, 2010], to calculate a p-value. We randomly shuffled the label (vectors) assignments for the EEG data such that the input EEG data *X* is randomly assigned to the labels *Y*. We refer to this randomized assignment as the shuffled dataset. This creates a situation where there is no association between the input *X* and the output *Y*. We then trained the prediction model on the shuffled dataset to obtain the permuted accuracy and repeated the shuffling procedure 100 times to obtain 100 permuted accuracies. We then fit a Gaussian kernel density estimator (KDE) function to the obtained permuted 2 vs 2 accuracies to generate a null distribution. The KDE has a smoothing effect on the distribution of the permutation accuracy values, allowing us to make inferences based on a finite number of permutation iterations. Finally, we compared the observed (non-permuted) accuracy value to the null distribution to obtain a p-value, and checked for statistical significance. To correct for multiple comparisons over time, we use the Benjamini-Hochberg-Yekutieli False Discovery Rate (FDR) correction method with no dependency assumption [Benjamini and Yekutieli, 2001].

### 2.8 Testing for Difference between Conditions

To test for significant differences in accuracy values between experimental conditions, we used the non-parametric cluster permutation test [Maris and Oostenveld, 2007]. We used an independent samples t-test (scipy.stats.ttest_ind) on the 2 vs 2 accuracy values for both age groups with 10000 permutations to obtain the p-values. Then, we used the Benjamini-Yekutieli procedure [Benjamini and Yekutieli, 2001] to correct the final p-values for multiple comparisons. Finally, we reported the largest cluster of time points containing at least 3 or more consecutive windows with p *<* 0.05.

### 2.9 Temporal Generalization Matrices

We used Temporal Generalization Matrices (TGM) [King and Dehaene, 2014] to assess the robustness of the neural representations across different time points. A TGM is obtained by training a model (as in Section 2.5) on data from one time window and testing the model on data from all the other time windows. The resulting matrix *M* contains cell values *M_ij_*, which denotes the accuracy values of the model trained on time window *i* and tested on time window *j*. If the neural representations in the beginning of the trial are similar to those later in the trial, the model performance will be similar across time windows.

## 3 Results

### 3.1 Analysis 1: Decoding Animacy Category from EEG

In this first analysis, we explored whether we can detect complex properties of nouns, such as animacy, in the infants’ neural representations. Here, we used a binary classification model to predict the animacy category of the stimulus word from the EEG data. Successful decoding will indicate that the neural patterns of the infants have representations of word animacy. We show the animacy decoding results in Fig 5. For 9-month-old infants, we observed significantly above chance accuracy at multiple time points, which shows that 9-month-old infants’ neural representations of single words include information about word animacy. Animacy representations appear to be strongest around 400ms and around 950ms after the onset of the word stimulus. This result is consistent with prior work showing that young infants have the ability to distinguish animate from inanimate objects [Rakison and Poulin-Dubois, 2001].

**Figure 5:**
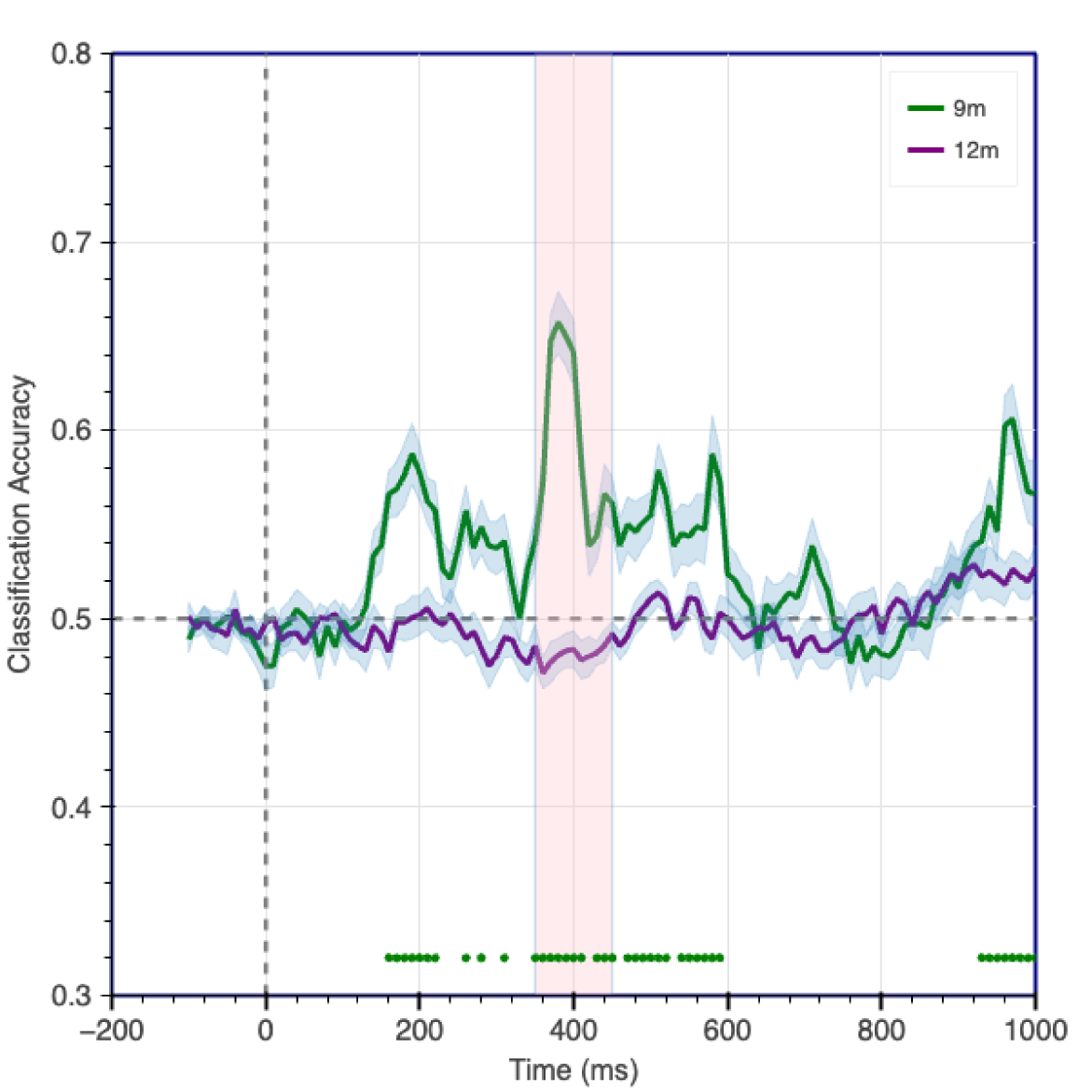
Accuracy for decoding word animacy category from the stimuli word. Green curve - 9-month-old infants; Purple curve - 12-month-old infants. Each point on the graph represents an accuracy value for the model trained on a 100ms window (100ms to the left of the accuracy point) with a 10ms sliding step. Green dots show significantly above chance accuracy points for 9-month-olds (p *<* 0.05, FDR corrected for multiple comparisons over time). No significantly above chance accuracy points were obtained for 12-month infants. The shaded (pink) area shows significant difference (p*<*0.05) between the accuracies of the two age groups using non-parametric statistical test [Maris and Oostenveld, 2007].

Interestingly, we could not predict word animacy with significantly above chance accuracy from the 12-month-olds’ EEG data. However, as we will observe in the next analysis, word semantic representations can be decoded from the neural data even though we did not find evidence of animacy decodability from the 12-month-old infants’ neural data.

### 3.2 Analysis 2: Representations of word semantics

It is established that young infants can understand the meaning of certain words from an early age. But how do the neural patterns change over time when engaged in the semantic processing of words? In this analysis, we explore this question. We used the prediction model (as in Section 2.5) to predict the word vectors for each stimulus word from the EEG data. We show the results in Fig 6.

**Figure 6:**
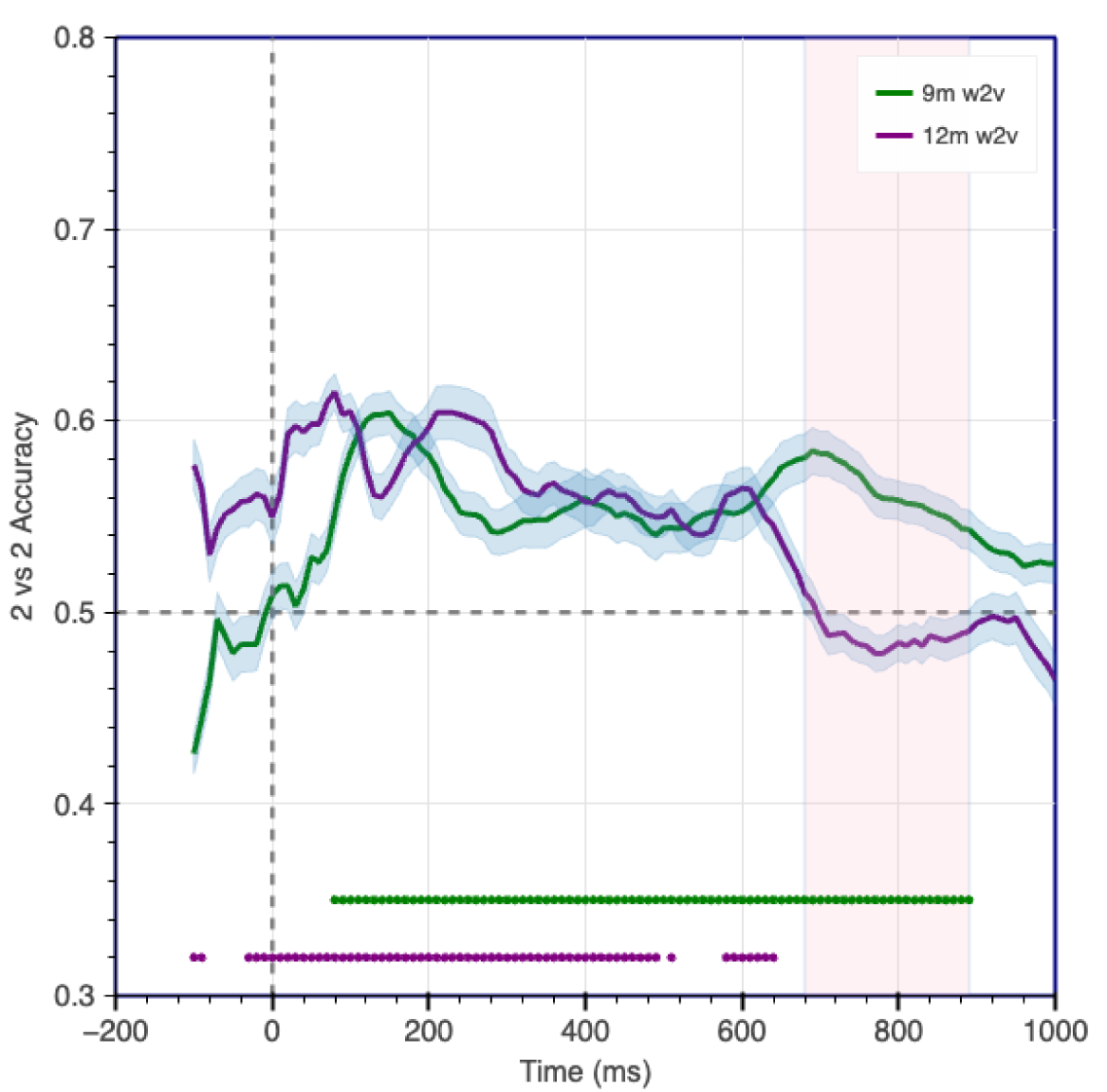
2 vs 2 accuracy for predicting original (pretrained) Word2Vec word vectors from EEG data collected from 9-month-old and 12-month-old infants. Each point on the graph represents an accuracy value for the model trained on a 100ms window (100ms to the left of the accuracy point) with a 10ms sliding step. The green dots (appearing as a continuous line) above the x-axis represent the points where the accuracy is reliable for 9-month-old infants, and the purple dots represent reliable above chance accuracy for 12-month-old infants (p *<* 0.05, FDR corrected for multiple comparisons over time).

We were able to predict the stimuli word vectors from the EEG data recorded from both 9-month-old and 12-month-old infants, which provides evidence for the presence of word semantic information in infant neural data at these ages. The results of this analysis also indicate that the infants’ neural representations of word semantics can be well characterized by the Word2Vec vectors of the stimuli words. In addition to existing work, which showed that infants can detect semantic mismatch of stimuli [Parise and Csibra, 2012, Friedrich and Friederici, 2010], our decoding analysis results show that the neural representations are also involved in processing single word semantics.

For 9-month-old infants, all accuracy values from the onset of the word until around 900ms after word onset were found to be above chance and statistically significant (green dots in Fig 6). For 12-month-old infants, we also observed above chance accuracy values for multiple time points after word onset but only until around 650 milliseconds, after which the accuracy dropped. The decoding accuracy also differed significantly late in the trial (pink shaded areas in Fig 6). This suggests a difference in the timing of the mental processes involved in the single word comprehension between the two groups of infants. ^4^

It must be noted that caution should be exercised while talking about the timing of such effects because the usage of a high-pass filter of 0.1Hz may have an effect on the timing of the signal onset. For example, we noticed a slight extension of the above chance accuracy into the baseline window, especially for the 12-month-old group. Such patterns have also been observed before in other decoding studies as well (e.g., Oh et al. [2019]). Thus, we took a number of steps to mitigate the effect. First, we ensured that all onset times for the stimulus words were manually checked for errors. Second, we explored other filtering bands as a value of 0.1Hz for the high-pass filter may have caused the spread of information across time and into the baseline window. Some previous work has suggested using other values for the high-pass component of the band pass filter, such as 0.5Hz [Choi et al., 2020, Gennari et al., 2021], and 0.3Hz [Debnath et al., 2020]. However, using a higher value for the high-pass filter resulted in poor model performance in both groups of infants compared to using a high-pass filter of 0.1Hz. Third, we also used robust detrending [van Driel et al., 2021] but it resulted in a reduction in model performance. All in all, we chose a band pass filter of 0.1-50Hz to ensure that important effect information is not discarded in the final analysis.

From this analysis, we observed that single word semantics can be decoded from the neural responses of infants. But such a result could be due to other confounding variables correlated with word semantics, which may have helped the model decode the stimuli. In the next analysis, we assess the presence of phonetic representations of word stimuli in the EEG data.

### 3.3 Analysis 3: Presence of phonetic representations of word stimuli

Although recent work has shown that phonetic features can be decoded from EEG data recorded from infants [Di Liberto et al., 2023], our data collection process preceded this finding. For our analyses, we selected the stimuli with the primary objective of decoding animacy categories of the stimuli, and given the limited vocabulary of infants this young, we did our best to balance for consonants, vowels, and syllable number. However, there is still a possibility of the phonemes being a confound to the semantic decodability of words. To this end, we carried out a simplified analysis where we observed the overall decodability of phonetic vector representation of words from the EEG data. Ultimately, we wanted to compare the semantic decoding curve and the phoneme decoding curve in order to observe significant differences. In other words, is there a presence of the semantic decodability at time points, where the phonetic decodability is absent? We aim to answer this question here, where we investigate the decodability of the phonetic representations of the stimuli words because it may be a possible confound to the decodability of the semantics, and to understand if there is a presence of semantic decodability in the absence of phonetic decodability.

To test the decodability of phonemes, we segmented each stimulus word into its individual phonemes. Each word in our stimuli set can be segmented into a combination of three phonemes to six phonemes. To obtain the individual IPA^5^ phonemes for the stimuli words, we used Canadian Pronunciation because the experimenter was a female native English Canadian speaker. When the Canadian Pronunciation was not available, we used the General American (GA) pronunciation. To obtain the pronunciations, we used the definitions in Dictionary.com^6^ and Wikitionary^7^. We mapped each phoneme to a 36-dimensional vector obtained developed previously [Mielke, 2012]. We concatenated all phonemes for a given word into one vector *P* and zero-padded all the concatenated vectors to make them equal in length, resulting in a 216-dimensional vector for each word. Finally, we trained a model as in Section 2.5 to predict the concatenated 216-dimensional vectors from the EEG data. We show the process of creating these concatenated phoneme vectors in Fig 7, and we show the results for decoding these concatenated phoneme vectors in Fig 8. For simplicity, we only explore the decodability of the overall representation of phonemes throughout the duration of the trial. Future work may look at the investigating the decodability of individual phonemes at different time points.

**Figure 7:**
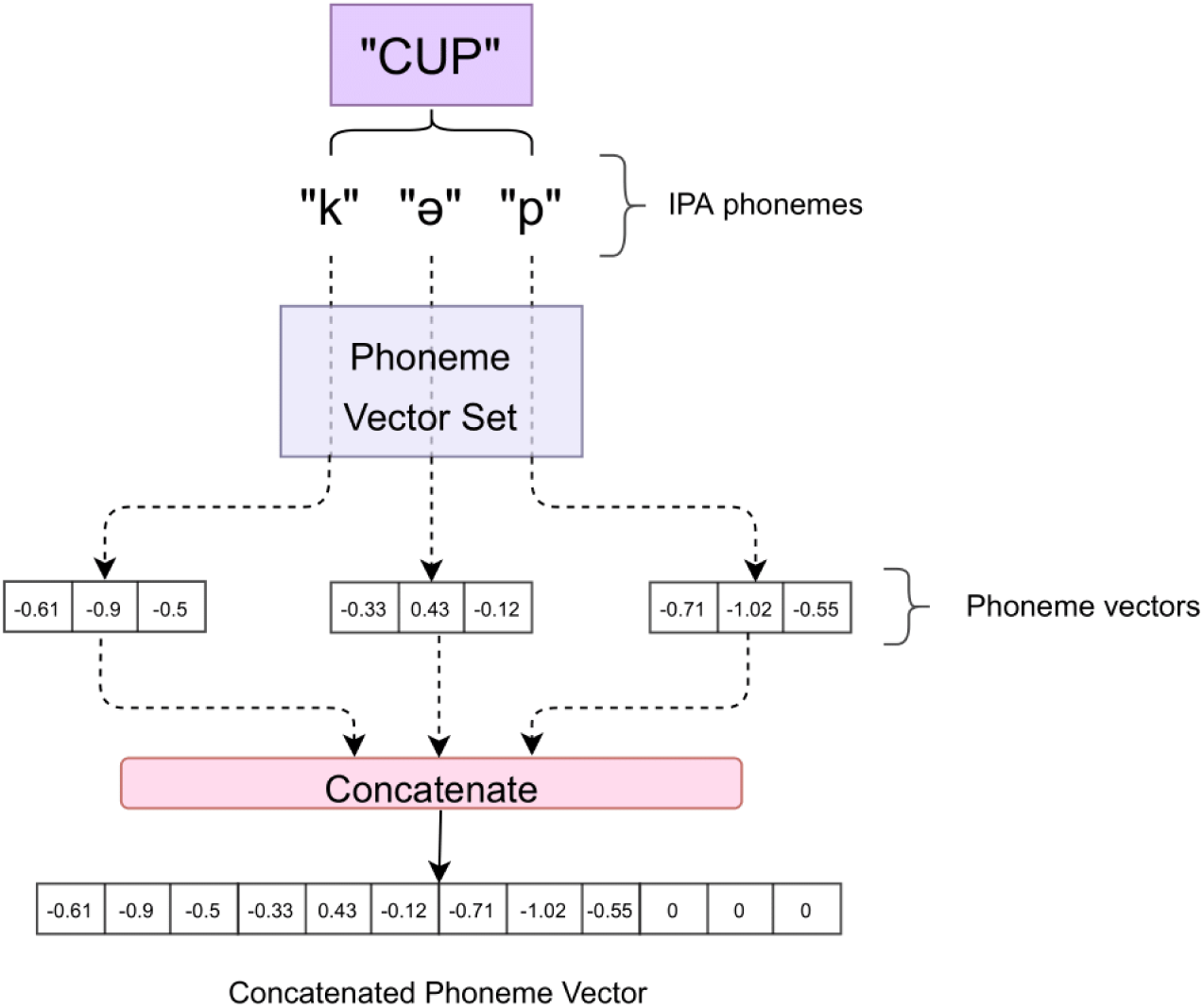
Phoneme vector creation process for an example word ’cup’ from the stimuli set. The stimulus word is broken down into its individual IPA phoneme. For each IPA phoneme, a vector is retrieved from a list of previously defined vectors [Mielke, 2012]. Finally, a vector is created by concatenating individual vectors. Zero-padding is used to obtain equal length vectors for all words. We show 3-dimensional phoneme vectors for simplicity. The vectors for individual phonemes are 36 dimensions long.

**Figure 8:**
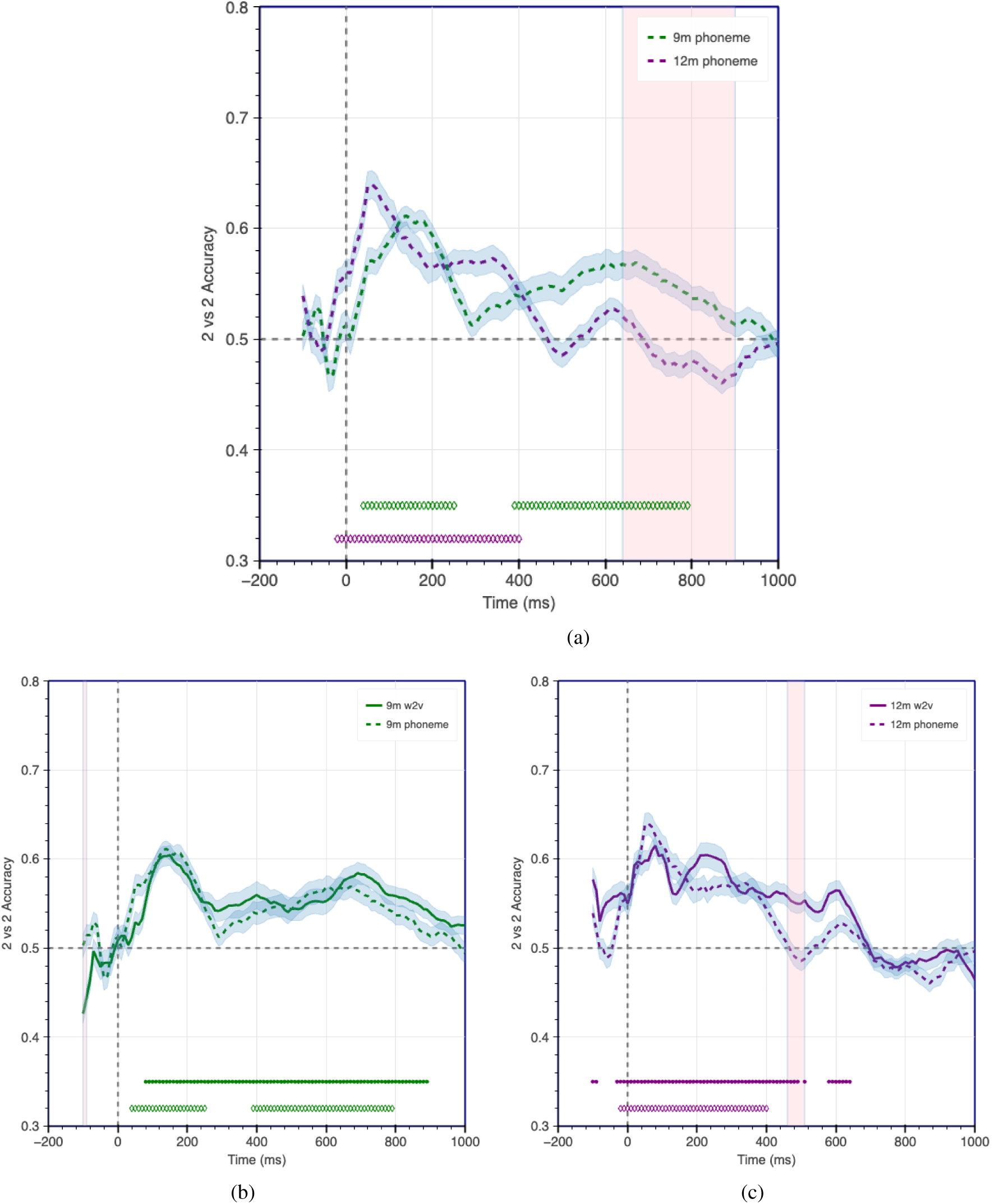
Decoding accuracy for predicting phonetic components of word stimuli. Fig 8a shows the decoding accuracy for predicting phonemes from EEG data recorded from 9-month-old and 12-month-old infants. Fig 8b shows the comparison of accuracy for predicting phonemes (green dashed curve) and original (pretrained) Word2Vec semantic vectors (green solid curve) from EEG data recorded from 9-month-old infants. Fig 8c shows the comparison of accuracy for predicting phonemes (purple dashed curve) and Word2Vec semantic vectors (purple solid curve) from EEG data recorded from 12-month-old infants. Shaded areas show significant differences in accuracies, dots above the x-axis show points of significantly above chance accuracy for decoding Word2Vec vectors, and ♢ symbols above the x-axis show points of significantly above chance accuracy for decoding phoneme vectors. p< 0.05, FDR corrected for multiple comparisons over time.

We were able to decode phonetic information with significantly above chance accuracy from the neural responses of 9-month-old infants (green dashed curve in Fig 8a). We observed significantly above chance accuracy immediately after word onset, which lasted till 800ms with a drop at around 300ms. On the other hand, the accuracy for the 12-month-old infants (purple dashed curve in Fig 8a) had an initial rise in decoding accuracy around word onset, which lasted till 400ms, quickly dropping after that. This drop suggests that though the 12-month-old infants encoded phonetic information, it may have been more transient than the phonetic representations for the 9-month-old infants. The pink shaded area starting about 650ms shows regions with significant differences between decoding accuracy for phoneme vectors for 9-month-old infants and 12-month-old infants.

We also compared the results between the current phoneme decoding analysis and the semantic decoding analysis described in Section 3.2, in Fig 8b for 9-month-old infants and in Fig 8c for 12-month-old infants. In Fig 8b, we see that for 9-month-old infants, the accuracy curve for the phoneme decoding analysis (green dashed curve) traces close to the accuracy curve for the semantic decoding analysis (green solid curve). When we compare the phoneme decoding analysis and semantic decoding analysis for 12-month-old infants, we see that in Fig 8c, the curve for phoneme decoding analysis (purple dashed curve) and semantic decoding analysis (purple solid curve) trace each other closely till around 400ms, after which the phoneme decoding accuracy drops sharply, but semantic decoding accuracy persists till about 650ms. This effect is depicted by the pink shaded area showing a significant difference in accuracy after 400ms. Such a result suggests that factors contributing to semantic understanding of words in 12-month-old infants after 400ms are not likely attributable to word phonetics but rather to other components of word semantics^8^. This result is different from that observed in the case of 9-month-old infants, where there was no significant difference between the phoneme and semantic decoding accuracy curves (no significantly noticeable pink shaded area). This finding suggests that we cannot rule out the possibility that the two curves are likely correlated to each other.

### 3.4 Analysis 4: Shared Representations of word semantics between age groups

In Section 3.2, we observed differences in both groups of infants with regard to the evolution of neural representations of word semantics. But are there underlying neural patterns that may be similar across age groups? To answer this question, we trained a ridge regression model on the EEG data from one group to learn the mapping between the EEG data and word vectors. Then, at test time, we used the EEG data obtained from the other age group to predict the word vectors. Successful decoding would indicate the presence of the shared neural representations of word semantics across age groups.

As seen in Fig 9, when we trained the model on EEG data recorded from 9-month-old infants and tested on EEG data from 12-month-old infants, we were able to reliably predict the word vectors (orange curve). We observed significantly above chance decodability from 250ms to 650ms. Similarly, when we trained the model on EEG data from 12-month-old infants and tested on EEG data from 9-month-old infants, we observed significantly above chance accuracy for the entire period from the beginning of the word onset to 850ms (black curve). The presence of above chance decodability when training and testing across age groups indicates that there exists similar neural representations of word semantics for both age groups. The decodability of word vectors from neural representations across age groups supports the idea that although there may be developmental changes across age groups, there is a significant similarity between them as well.

**Figure 9:**
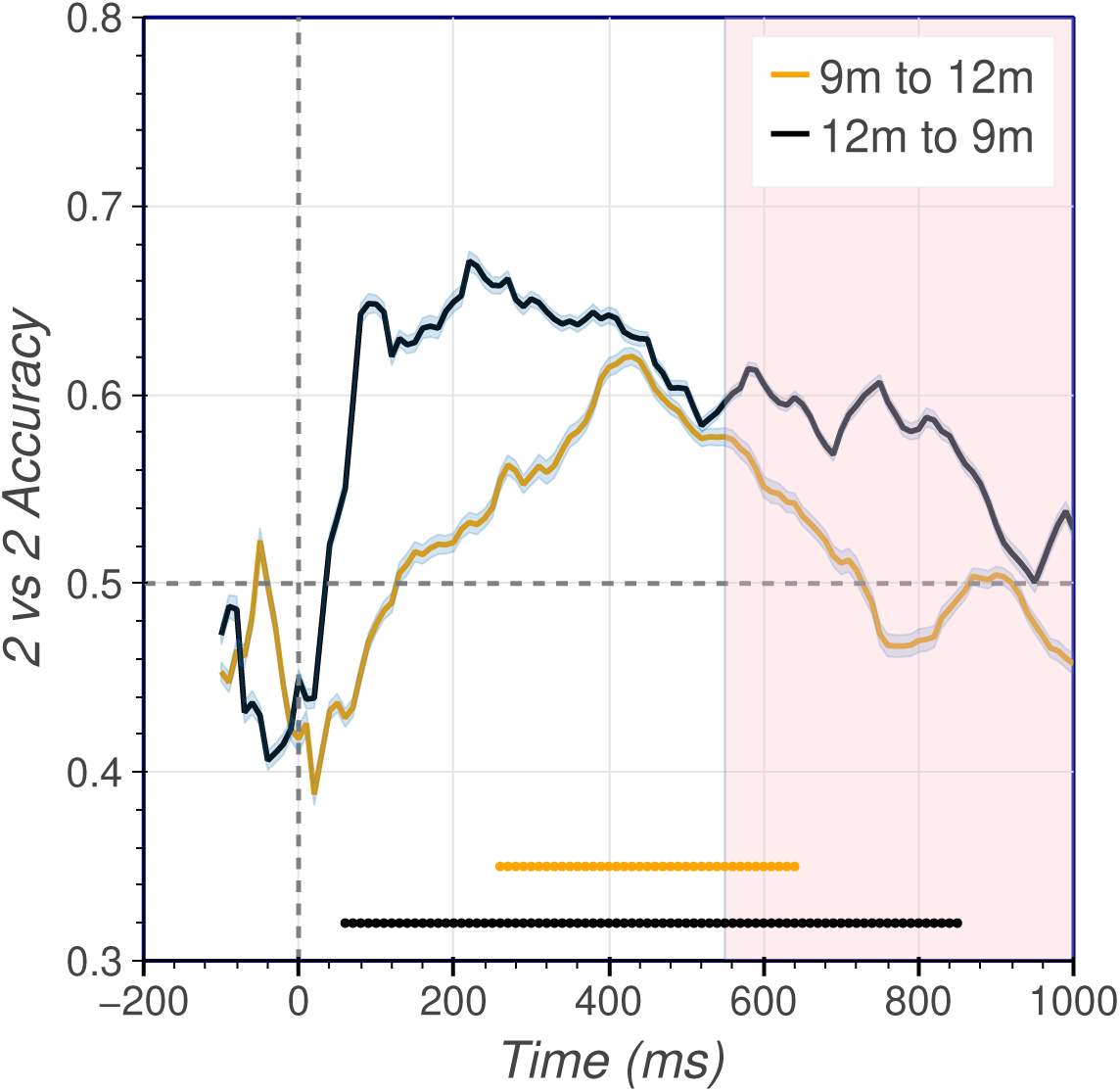
2 vs 2 accuracy for predicting Word2Vec word vectors from EEG across age groups. Each point on the graph represents an accuracy value for the model trained on a 100ms window (100ms on the left of the accuracy point) with a 10ms sliding step. The orange dots denote above chance accuracy for the model trained on 9-month-old and tested on 12-month-old infants, and black dots denote above chance accuracy for the model trained on 12-month-old infants and tested on 9-month-old infants (p *<* 0.05, FDR corrected for multiple comparisons over time). The shaded area shows where the accuracy curve is significantly different for the two conditions.

### 3.5 Analysis 5: Decoding fine-tuned word vectors from EEG

In Section 3.2, we predicted the Word2Vec vectors from EEG. To obtain these vectors, the Word2Vec model was trained on the Google News Corpus. The resulting word vectors will likely better represent adults’ understanding of word meaning because the news articles are written for adults. It may be possible that the pretrained word vectors are not ideal representations of a child’s understanding of word meaning. Word vector representations that better represent how infants understand word meaning may lead to better decodability than the original (pretrained) Word2Vec vectors. To this end, we fine-tuned the Word2Vec model on two datasets, the children’s book test dataset [Hill et al., 2016] and the Childes corpus dataset [MacWhinney, 2000]. For the fine-tuning process, we used one thousand iterations of gradient descent with default model hyperparameters^9^ (start learning rate: start_alpha=0.025, end learning rate: end_alpha=0.0001). We then train a similar prediction model (as in Section 2.5) to decode the fine-tuned word vectors.

Fig 10a shows that the fine-tuned word vectors can be decoded with significantly above chance accuracy from the EEG data from both infant groups. However, there is no significant difference when comparing the decoding accuracy of the original pretrained vectors and the decoding accuracy of the fine-tuned word vectors. In Fig 10b, we notice that the decoding accuracy for the 9-month-old infants for the original word vectors (green solid curve) is similar in time course compared to the decoding accuracy for the fine-tuned word vectors (green dotted curve). Also, for the 12-month-old infants, we observe similar results in Fig 10c, where the decoding accuracy for pretrained word vectors (purple solid curve) follows closely to the decoding accuracy of fine-tuned word vectors (purple dotted curve). Since using fine-tuned vectors did not perform better than their pretrained counterpart, we used pretrained word vectors for our analyses.

**Figure 10:**
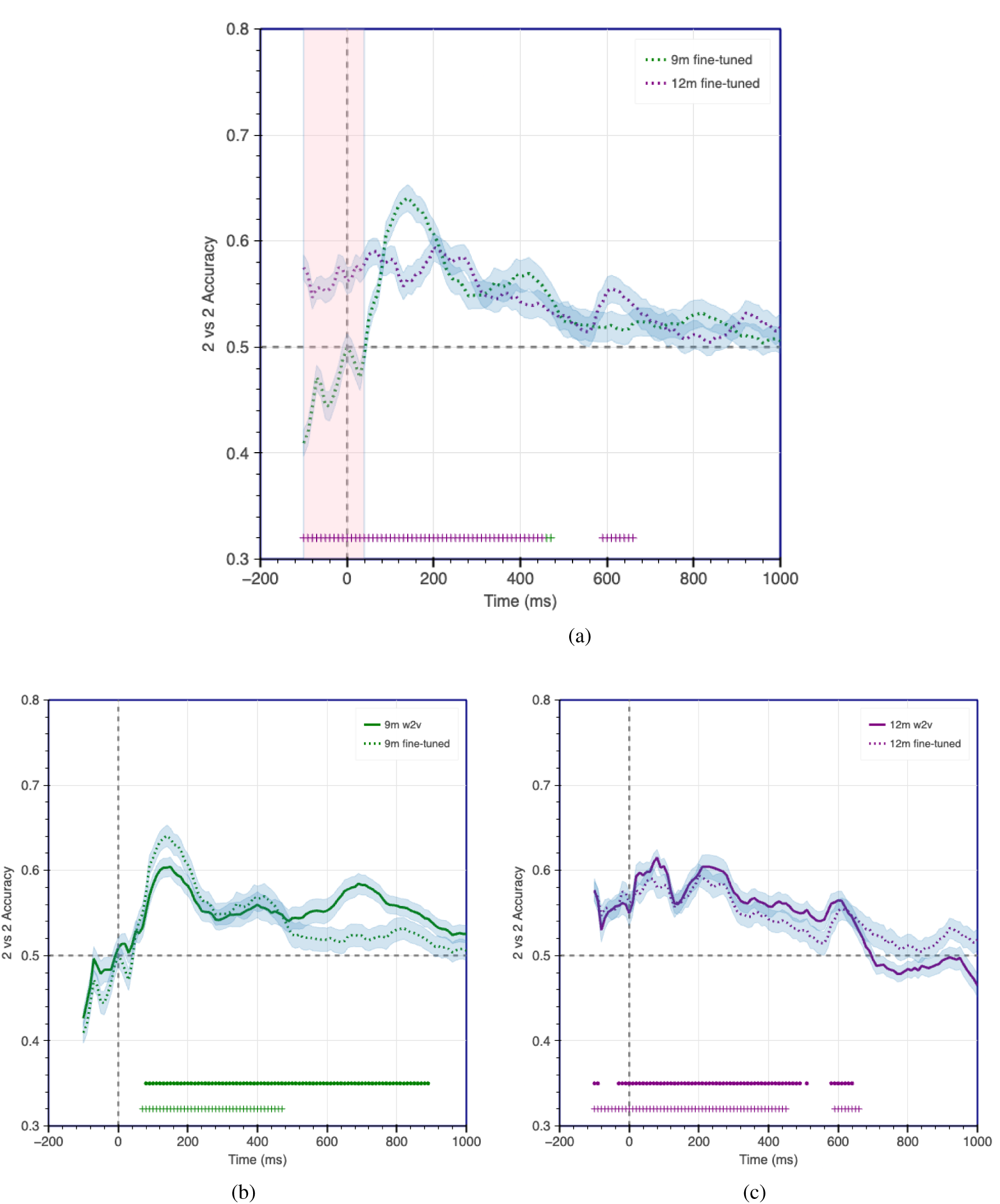
10a shows the decoding accuracy for predicting fine-tuned word vectors from EEG data. 10b shows the accuracy comparison for predicting fine-tuned word vectors (dotted green curve) and original pretrained word vectors (green solid curve) from EEG data collected from 9-month-old infants. 10c shows the accuracy comparison for predicting fine-tuned word vectors (purple dotted curve) and pretrained word vectors (purple solid curve) from EEG data collected from 12-month-old infants. Shaded areas show significant differences in accuracies, dots above the x-axis show points of significantly above chance accuracy, and the + symbols show points of significantly above chance accuracy for decoding fine-tuned word vectors. p*<* 0.05, FDR corrected for multiple comparisons over time.

### 3.6 Analysis 6: Generalizability of neural representations for word semantics

In the previous analyses, we investigated the emergence of the detectable neural representations of word meaning as the infants listened to single words. The question then arises, do these neural representations generalize across time? In other words, do the neural representations from one time window reemerge later during the trial, and are these representations sustained or evolving over time? To answer this question, we used Temporal Generalization Matrices (see Section 2.9). The neural representations of word semantics will be similar between two time windows if the model trained on one time window (see Section 2.5) is able to decode the word vectors from a different time window. We trained and tested the model on all the possible pairs of time windows to obtain a matrix of 2 vs 2 accuracies. We show the TGM results in Fig 11 where the significantly above chance accuracies are indicated by black bounding contours.

**Figure 11:**
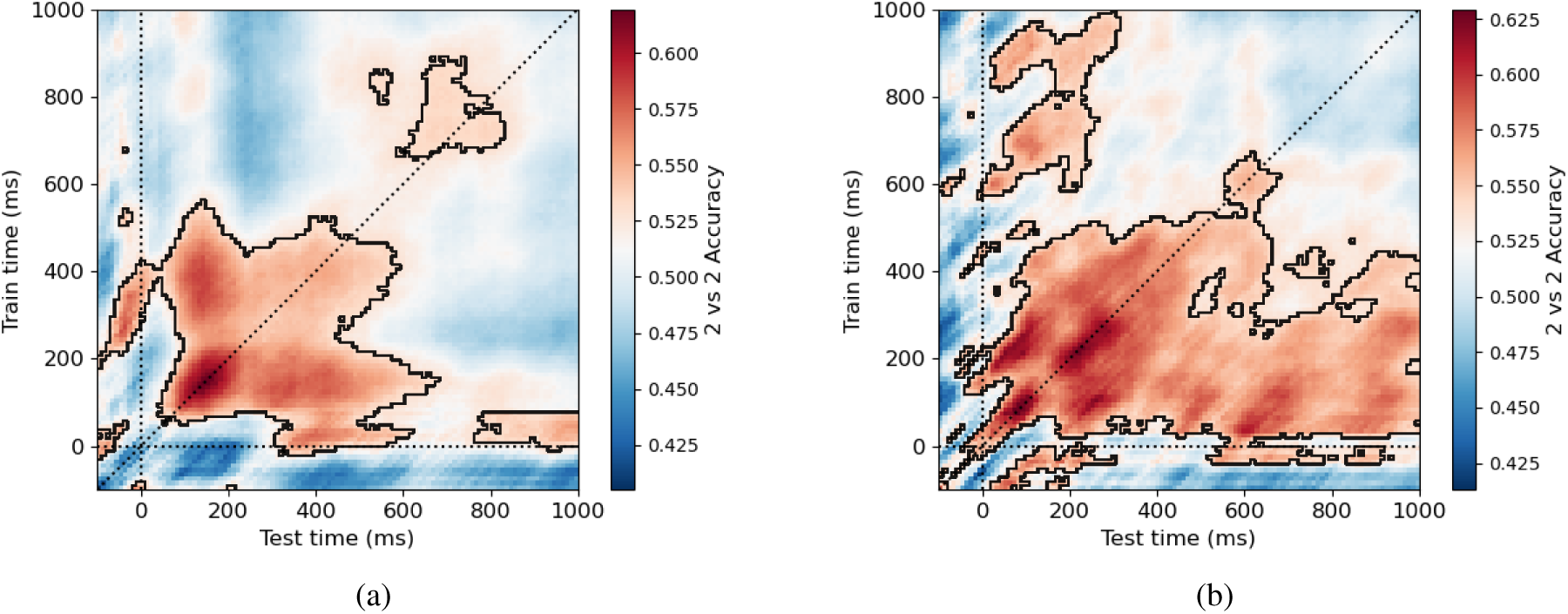
Temporal Generalization Matrices for 9-month (a) and 12-month-old infants (b). To obtain the matrix, we trained the regression model on EEG data from one time window and tested on all possible time windows. Fig 11a shows the temporal generalizability of neural representations of word semantics in 9-month-old infants. There is a weak generalizability, showing patterns of neural activation in the beginning of the trial and during the end of the trial. 11b However, for 12-month-old infants, the neural patterns show stronger generalizability as observed by the significantly above chance accuracy areas further away from the diagonal.

Fig 11a, shows the TGM decoding results for the 9-month-old infants’ neural data. We observe statistically significant results mostly immediately after word onset, along the diagonal, with areas extending away from the diagonal. The extension of significantly above chance results away from the diagonal indicates generalizability of the neural representations The neural representations of word semantics in the beginning of the trial (50ms after word onset) are similar to those observed for the whole duration until before 600ms. The above chance accuracy starting later in the trial (around 750ms) along the diagonal may suggest the presence of isolated mental processes in 9-month-old infants related to the comprehension of single word semantics.

Fig 11b shows the TGM decoding results for the 12-month-old infants’ neural data. In comparison to the 9-month-old infants, the 12-month-old infants’ off-diagonal accuracy extends from the beginning of the trial (right after word onset) till the end of the trial (around 1000ms). Moreover, there were more off-diagonal areas of significantly above chance accuracy compared to those observed for the 9-month-old infants. An interesting observation is that of the presence of off-diagonal accuracy late into the trial (from 700ms-1000ms) without a strong presence of diagonal accuracy (from 700ms-1000ms). This may suggest that there may be other noise characteristics contributing to such a phenomenon. All in all, these patterns suggest that the neural representations of word semantics for the 12-month-old infants are more stable and generalizable than those of the 9-month-old infants.

### 3.7 Analysis 7: Generalizability of neural representations across age groups

Our final question is that of generalizability of word semantic representations across time and age groups. We ran the Analysis in Section 3.6 again, but this time evaluated the model using the data from the other age group. We show the results in Fig 12.

**Figure 12:**
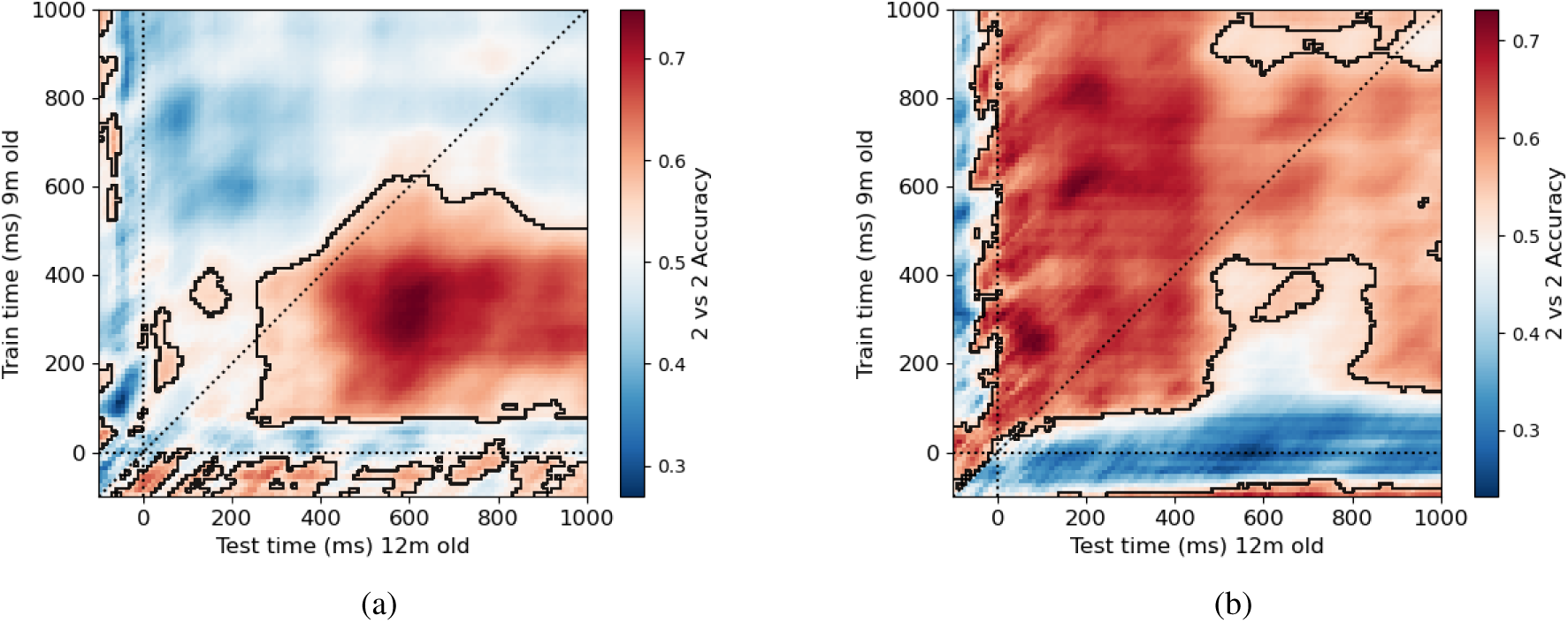
Temporal Generalization Matrices for training and testing across age groups. To obtain the matrix, we trained the regression model on EEG data from one time window and tested the model on all the possible time windows. Fig 12a shows the temporal generalizability of neural representations of word semantics when the model was trained on neural data recorded from 9-month-old infants and evaluated on neural data recorded from 12-month-old infants. We observe that neural representations present early during the trial for 9-month-old infants show similarity to those occurring later in the trial for 12-month-old infants suggesting similar mental processes are involved in word comprehension, but at different time points. Similarly, in Fig 12b, we observe areas of significantly above chance accuracy, mostly above the diagonal, indicating that majority of the neural representations for 12-month-old infants involved in word comprehension are similar to neural representations occurring early during the trial for 9-month-old infants.

When we train our model on the EEG data recorded from 9-month-old infants and evaluate it on EEG data recorded from 12-month-old infants (Fig 12a), a large area of significantly above chance accuracy is observed on the off-diagonal location. More specifically, training our model on early windows of 9-month-olds’ EEG data (around 100ms to 600ms) shows significantly above chance accuracy for the parts of the trial observed slightly later (from 300ms to 1000ms). This indicates that representations of word semantics present in 9-month-old infants are similar to those present in later time points in the trial for 12-month-old infants. Fig 12b supports this result. We also observed areas of above chance accuracy primarily above the diagonal, indicating that the representations identified by the model from the EEG data from 12-month-olds are similar to representations identified early during the trial from 9-month-old infants across multiple time points.

### 3.8 Analysis 8: Electrode level decoding

In Section 3.2, we observed the semantic decoding accuracy for both groups of participants. In addition, we were also interested to understand the regions of activation of the neural representations in the two groups of participants. To this end, we ran the same analysis in Section 3.2 on the channel level, where we trained the prediction model on the groups of channels (by considering a channel and its immediate neighbours to reduce noise) to predict the semantic vectors. First, we chunked the EEG data into 300ms windows (in contrast to 100ms in previous experiments). For each window, we considered the data from each electrode (and its immediate neighbours to reduce noise), on which we trained the prediction model (from Section 2.5). then for each window, we aggregated the results and use mne ^10^ to plot the topomaps. We used the windows 0-300ms, 300-600ms, and 600-900ms to display the results. We chose the 0-300ms window to observe the activations of the EEG data during the peaks of the decoding accuracy(see Fig 6 in Section 3.2). The 300-600ms window allowed us to observe the activation patterns during the times when semantic decoding accuracy is relatively similar for both groups of participants. Finally, choosing a 600-900ms window showed us the areas of activation during time periods of significant differences in decoding accuracy (pink shaded area in Fig 6). We show the activated areas for the 2 vs 2 accuracy in Fig 13.

**Figure 13:**
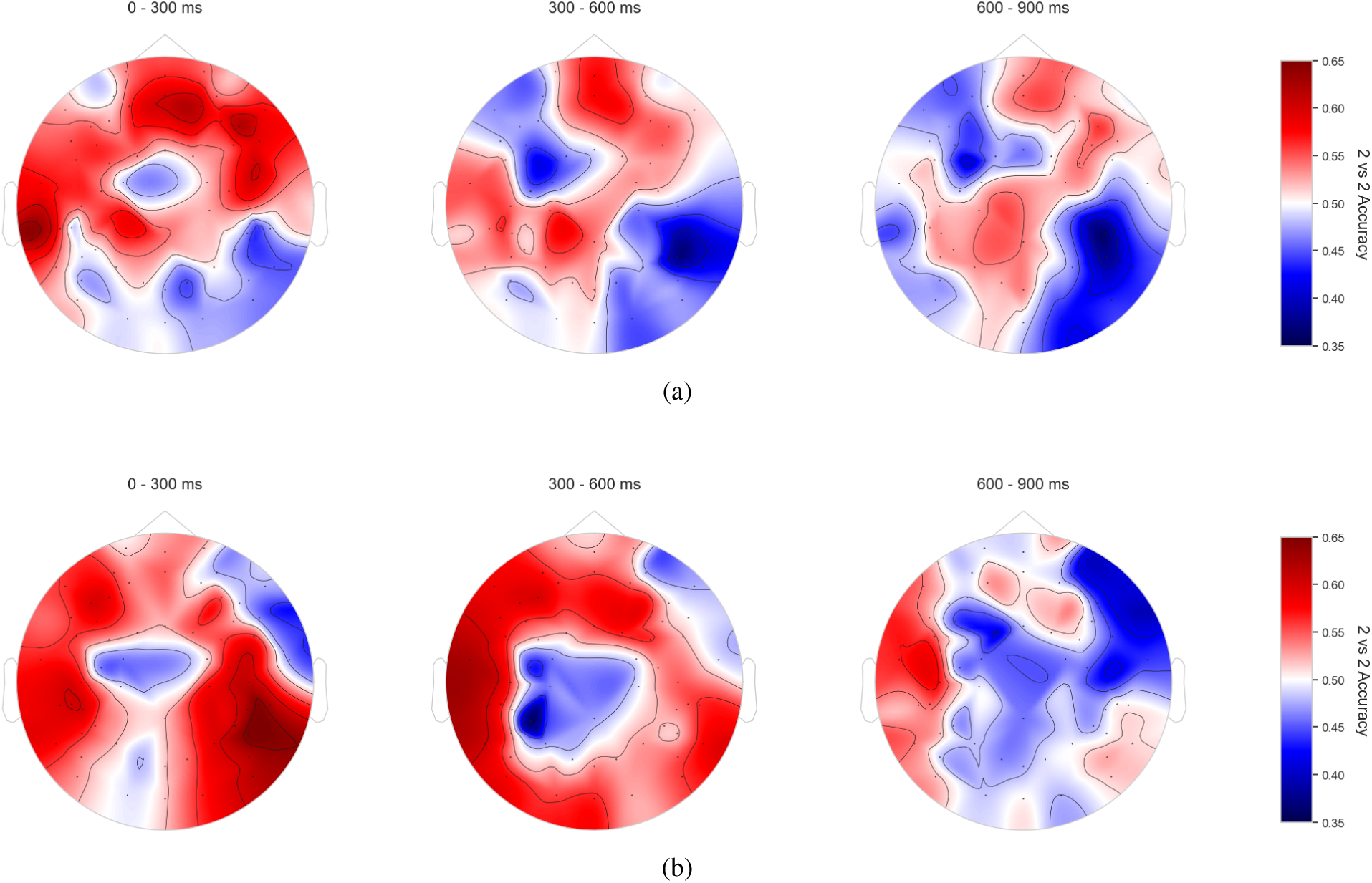
Topomaps for 9-month-old (a) and 12-month-old (b) participant groups. Activation patterns appear to be relatively distributed, with concentrated areas of higher 2 vs 2 accuracy during the 0-300ms and 300-600ms windows. The windows of 0-300ms and 300-600ms windows appear to be most different for both groups of participants.

Generally, the activations appear to be distributed over the entire brain region in both groups of participants. For both the 9-month-old and the 12-month-old group, we observed multiple regions of high accuracy during the 0-300ms and 300-600ms windows. During the 600-900ms window, we observed relatively lower 2 vs 2 accuracy values for the localization patterns relative to the earlier windows. Moreover, activation patterns are the most different between the 0-300ms and the 300-600ms windows for both groups of participants. Such a result may indicate a change in the mental processes involved in processing word semantics, from the 0-300ms to the 300-600ms window.

## 4 Discussion

In this work, we explored the dynamics of neural representations in infants while processing single word semantics. To this end, we used a decoding approach on EEG data obtained from 9- and 12-month-old infants. Our results indicate that the neural representations engaged in the processing of single word meaning are rapidly activated after the onset of the word, with a statistically significant above chance decodability in both groups of infants. Notably, the neural representations for 9-month-old infants persist for a duration longer than those of the 12-month-old infants. We further observed that the decoding accuracy predicting word semantics may be correlated to that of word phonetics in both infant groups. Despite developmental changes between the 9- and 12-month-old groups, we also found shared aspects of neural representations of word semantics for both age groups. Furthermore, our results demonstrate that the neural representations of word semantics generalize across different time windows and age groups, as observed in the Temporal Generalization Matrices.

To the best of our knowledge, ours is the first work that investigates how neural representations in infants change over time during the processing of single word meaning. Previous research indicates that preverbal infants start to categorize objects into taxonomies [Behl-Chadha, 1996] but are also sensitive to both featural and thematic relations [Peters and Borovsky, 2019]. Infants also have the capacity to distinguish between animate and inanimate objects, which may be an important developmental foundation for understanding the world around them [Rakison and Poulin-Dubois, 2001], and arguably also for building representations of word meaning. As a first step towards decoding complex word properties such as word semantics, we began by decoding the animacy category of the word in the context of understanding word meaning. Our results show that for 9-month-old infants, the animacy category of the stimuli words can be decoded with significantly above chance accuracy around 400ms post word onset. This is in agreement with previous ERP studies supporting the role of the N400 signal in word semantics comprehension [Junge et al., 2012, Parise and Csibra, 2012, Cosper et al., 2020]. In the case of 12-month-old infants, we did not observe significantly above chance animacy decodability. It may be possible that the infants in the 12-month-old group exhibited weak signal of animacy. However, we were still able to observe significantly above chance word semantic decodability as we discuss next.

While decoding semantics representations of words from the neural data, both groups of infants showed prolonged decodability after word onset. The decoding accuracy for the 9-month-old infants was significantly above chance till the end of the trial. However, the decoding accuracy for 12-month-old infants dropped to chance levels after 650ms of word onset, suggesting that the neural representations of word semantics for the 12-month-old infants are not as long lasting as those of the 9-month-old infants. A possible explanation for such an effect is that older infants may be processing information faster than younger infants because their language processing capacity is more advanced. In fact, there is evidence in previous ERP studies indicating that as infants grow older, the timing of the N400 signal onset decreases and approaches closer to 400ms. However, there does not appear to be a significant difference in the onset of the N400 between 9- and 12-month-old infants (see [Junge et al., 2021] for a review). Another possible explanation is that older infants complete semantic processing of the word faster than younger infants. Our work is a first step towards connecting previous studies using ERP and decoding techniques to study word semantic comprehension in infants.

We observed in Analysis 2 (Section 3.2) that the peak decoding accuracy value was similar for both groups of infants. This suggests that the neural representations of both age groups are almost equally capable of representing single word meaning. The accuracy peak is observed immediately after word onset, meaning that the representations appear to be the strongest during the early part of the trial. This result is in agreement with previous work inspecting word understanding with looking time paradigms [Bergelson and Swingley, 2012]; even though it takes some time for infants to initiate eye movements in response to a speech signal, the neural representations for word meaning emerge immediately after word onset.

As discussed in Section 3.3, the infants can discriminate the phonetic features of words from early in life, and that over the first year of life, this ability becomes increasingly attuned to the phonetic differences used in their native language [Werker and Tees, 1984, 1992, Jusczyk and Aslin, 1995, Cheour et al., 1998]. The phonetic representations of infants over the age of 1 year have also been studied through the lens of looking time paradigms [Swingley and Aslin, 2000, Mani and Plunkett, 2008], which show differences in the infants’ ability to recognize the pictures when the stimulus word is mispronounced. A recent study has also shown that the envelope of audio stimuli can be reconstructed using the EEG data recorded from infants, further suggesting that infants have strong neural signals related to processing rhythmic patterns of speech [Attaheri et al., 2022], which may help them to understand semantics. However, there has been less work on the extent to which individual phonetic information of word stimuli is represented in the lexicon of infants younger than one year. Some previous research has shown that individual speech tokens such as vowels and consonants can be accurately classified from EEG data recorded from infants as young as 3-months-old [Gennari et al., 2021], and provides evidence for the ability to discriminate individual linguistic features in young infants. Nonetheless, our analysis (in Section 3.3) provides insights into how phonetic information is processed at a word level in conjunction to understanding word semantics. In the case of the 12-month-old group of infants, the phoneme decoding accuracy curve also closely followed the word semantics decoding accuracy, but only until around 400ms, after which the phoneme decoding accuracy fell to chance levels. This suggests that the neural representations after 400ms engaged in word semantic comprehension may not be confounded by the word phonetics, and we can be more certain it is directly related to word semantics. This result opens up new avenues of research for further exploring how neural representations of word semantics may be related to other components of word meaning.

As infants grow older, numerous factors, such as social interactions, exposure to more words, and other developmental changes, can collectively aid in the changing of the neural representations of word semantics. But do the neural representations engaged in understanding single word semantics for the older infants differ from those of the younger infants? Our results suggest that even though brains mature as infants grow older and infant vocabularies become richer, the neural representations involved in understanding single word semantics share similarities. In Section 3.4, we were able to decode the word semantic representations when the model was trained and tested across age groups. The decoding patterns, however, were different when training on data from 9-month-old infants versus training on data from 12-month-old infants. This difference could be due to less consistent data in the younger infant group, possibly stemming from less developed and more variable word representations as compared to the 12-month-old infants. Prior work has found that infants in the age group of 11-14 months old have adult-like characteristics in understanding word meaning [Bergelson and Swingley, 2018], which may further speak to the idea that older infants have more consistent neural representations of word semantics than younger infants. Such ideas may be explored in future work.

In language processing, the generalizability over time of neural representations in adults has been studied using various modalities [Fyshe et al., 2019, Honari-Jahromi et al., 2021], but there is a lack of work studying such generalizability in infants. Our results shown in Sections 3.6 and 3.7 are the first of their kind to analyze the generalizability of the neural representations of word semantics in infants. The neural representations of word semantics were generalizable across time windows and across age groups (see Figs 11 and 12). The neural representations of word semantics seemed to be more generalizable for 12-month-old infants than those of 9-month-old infants. The 9-month-old infants also showed early signs of generalizability. These results may suggest that as infants grow older, their neural representations of word semantics become more stable, possibly due to their exposure to more vocabulary and social experiences.

From Section 3.4, we observed the presence of shared aspects of word semantics in the neural representations for both age groups. In other words, the developmental changes across age groups may not necessarily lead to a complete change in how the neural patterns of infants represent word meaning. We see in Section 3.7, when testing for generalizability across age groups, there were large areas of significantly above chance accuracy. When training our model on the 9-month-old infants’ data and testing on the 12-month-old infants’ data (Fig 12a), we observed a large area of significantly above chance decodability. The area of significantly above chance decodability was larger when we trained our model on the 12-month-old infants’ data and tested on the 9-month-old infants’ data (Fig 12b). Such a result may suggest that as infants grow older, the neural representations of word semantics build on top of those formed earlier in life. It must be pointed out that a longitudinal study would help us understand the likelihood of such an effect more clearly, which we leave for future research.

We also wanted to explore the possibility of further improving the quality of the word vector representations of word semantics in order to increase the decoding accuracy. In Section 3.5, we fine-tuned the Word2Vec model on textual data that may be more representative of the infant lexicon. Although we hoped that fine-tuning Word2Vec would lead to better decoding accuracy, we did not observe this effect. It is possible that the child-directed speech data is too small to conduct a modelling experiment. It is also possible that by using speech directed at older children rather than exclusively infants, we introduced additional noise that further mismatched the word vectors to the infants’ lexicon. Fine-tuning is a well-explored strategy to obtain task-specific word vector representations, but more research is needed to study the techniques of fine-tuning word vectors for decoding word semantic representations in infants.

Finally, our decoding analysis on the electrode level provided insights into the distribution of the neural activations engaged in processing word semantics (Section 3.8). From our analysis in the 9-month-old group, we observed activations primarily over the frontal areas during the 0-300ms part of the trial. Such results are consistent with those reported in previous work conducting an N400 analysis (see Junge et al. [2021] for a review), where the most relevant brain regions of 6-11 months old were along the right, right-midline, central, central-parietal and the parietal regions of the brain. We also observed activations from electrodes in the left hemisphere of the brain in the 0-300ms and the 300-600ms part of the trial, which may further suggest signs of involvement of the language areas of the brain in processing single words semantics at an early age.

For the 12-month-old group, during the 0-300ms window, we observed activations from electrodes in both the left and the right hemisphere, and activations from the electrodes in the left hemisphere in the 300-600ms window. This may suggest that activations are likely localized to the left hemisphere of the brain during the middle of the trial. Later into the trial (600-900ms), the strength of the representations appear to reduce, with slight dominance of activations from the electrodes in the left hemisphere. This is in contrast to activated areas observed in the 9-month-old group, where during the middle (300-600ms) and later parts of the trial (600-900ms), the activations appeared to be more distributed. Such a result may further hint at the developmental changes that occur with respect to language understanding. In other words, as infants develop, the cognitive processes involving word semantic understanding become more left lateralized. A longitudinal study will further provide insights into how developmental changes occur at a participant level, which we leave as an idea for future research.

All in all, our electrode level analysis shows the activations of the neural representations pertaining to processing single word meaning, how these activations differ across the two groups of participants, and how these activations change over the course of the trial. These results can serve as a foundation for studying the changes that occur in the brain during the early years of language acquisition.

## 5 Conclusion

In this work, we studied the dynamics of the neural representations of word semantics in young infants using a decoding approach. Our analysis showed that single word semantics can be predicted from the neural data immediately after word onset, which extended until later in the trial. The entirety of the neural representations of word semantics for 9-month-old infants may be correlated to their ability to distinguish phonemes. However, for 12-month-old infants, this appeared to be the case only until 400ms indicating that neural representations of word semantics after this time period may correspond to other aspects of word semantics. Despite there being developmental changes between the two groups of infants, the neural representations involved in word semantics comprehension also shared similar characteristics. Moreover, our TGM analysis showed early signs of generalizability of the neural representations not only across time windows but also across age groups.

Our work raises several new questions. We observed that the decoding accuracy for word semantics for the 12-month-old infants falls to chance levels quicker than those of 9-month-old infants (Section 3.2). What is the cause of such an effect? Do the older infants process word semantics faster than the younger infants? Further research is required to disentangle this question. Although it is possible that the majority of the word semantics comprehension highly corresponds to the ability to distinguish phonemes, it is unclear as to what aspects of word semantics can be attributed to the above chance decodability of word semantics for 12-month-old infants after 400ms (Fig 8c), and more research investigating various properties of word semantics is required to answer this question. Our work uses word vector representations from the Word2Vec model. We note that more recent work with adults has used representations from the intermediate layers of advanced neural network models such as Convolutional Neural Networks and Transformers, to show that the representations generated in such models are similar to those in the adult brain [Caucheteux and King, 2022]. It may be illuminating in future work to explore the correlation between the neural representations of infants and the hidden representations of such advanced neural networks. Finally, it will be revealing to compare the neural representations of single word semantics in infants to those of the adults. The similarities and differences seen in the neural representations will provide insights into how semantic representations change as infants become increasingly sophisticated language users.

## Data and Code Availability

The code for this work is available at https://github.com/simpleParadox/eeg_semantics. The raw and preprocessed data is available at https://osf.io/hca7b/.

## Acknowledgements

We thank Jeff Mielke for providing us with the vector representations of the phonemes and for answering our questions.

## Supplementary Materials

**Figure S1:**
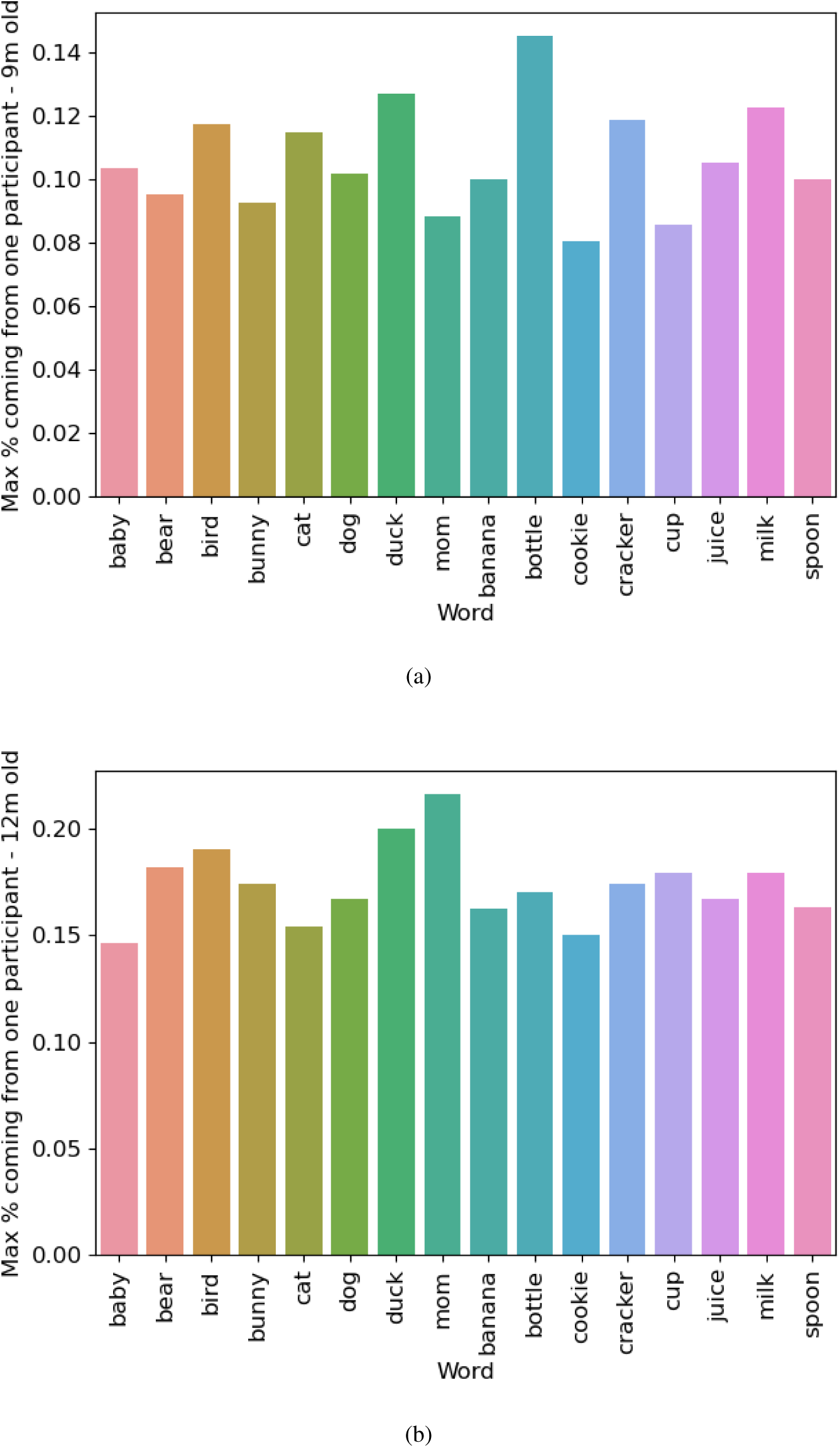
Maximum percentage contribution of each word to the whole dataset coming from one participant in the 9-month-old group (a) and the 12-month-old group (b). The word ’bottle’ is the word with the highest percentage coming from one participant in the 9-month-old group (14.5% of all trials). From the 12-month-old group, the word is ’mom’ (21.6% of the total trials for ’mom’) in the 12-month-old group.

**Figure S2:**
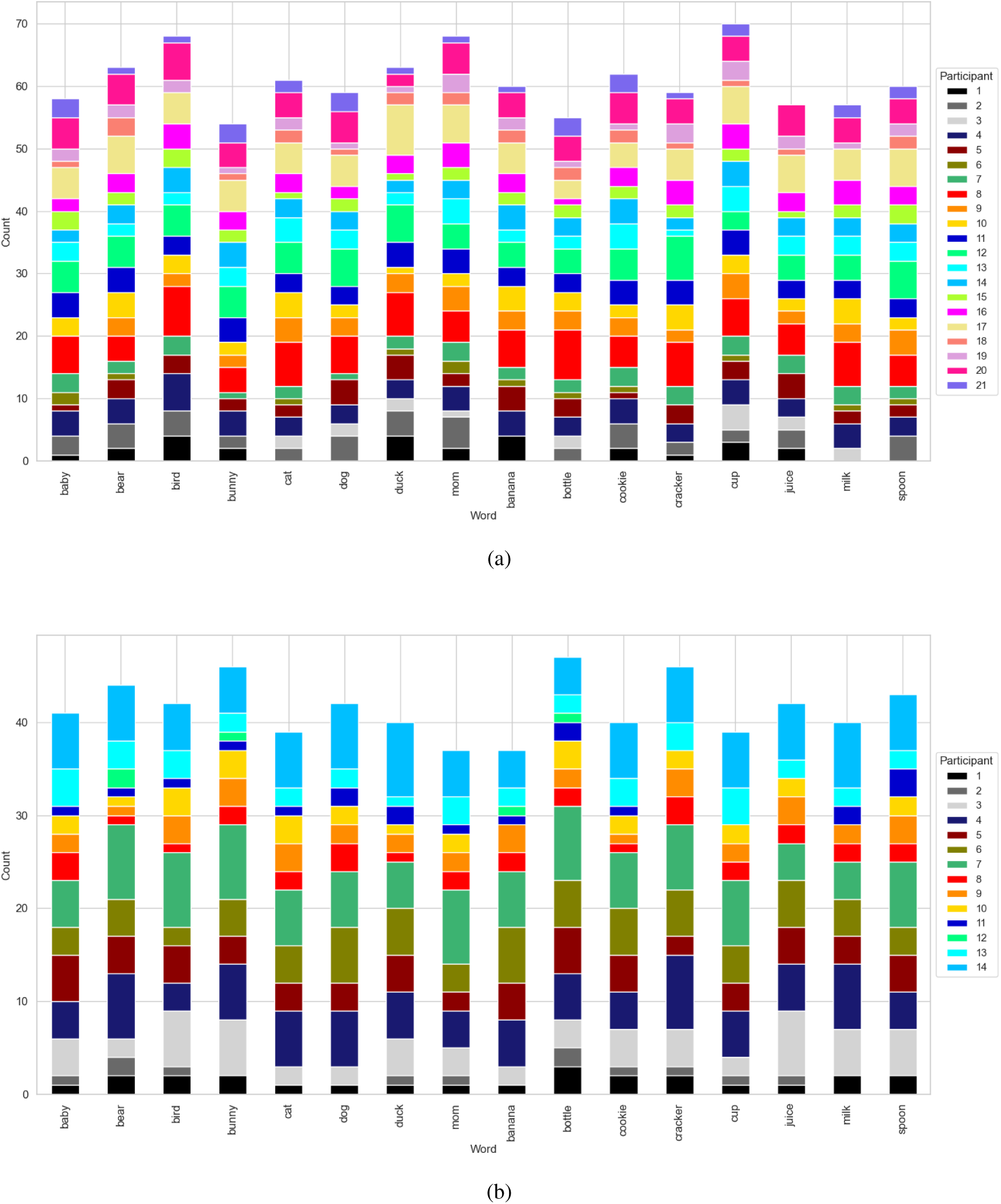
Number of good trials for each word stimulus per participant in the 9-month-old (a) and 12-month-old (b) participant group. X-axis denotes the word, and the y-axis denotes the count for each participant.

**Figure S3:**
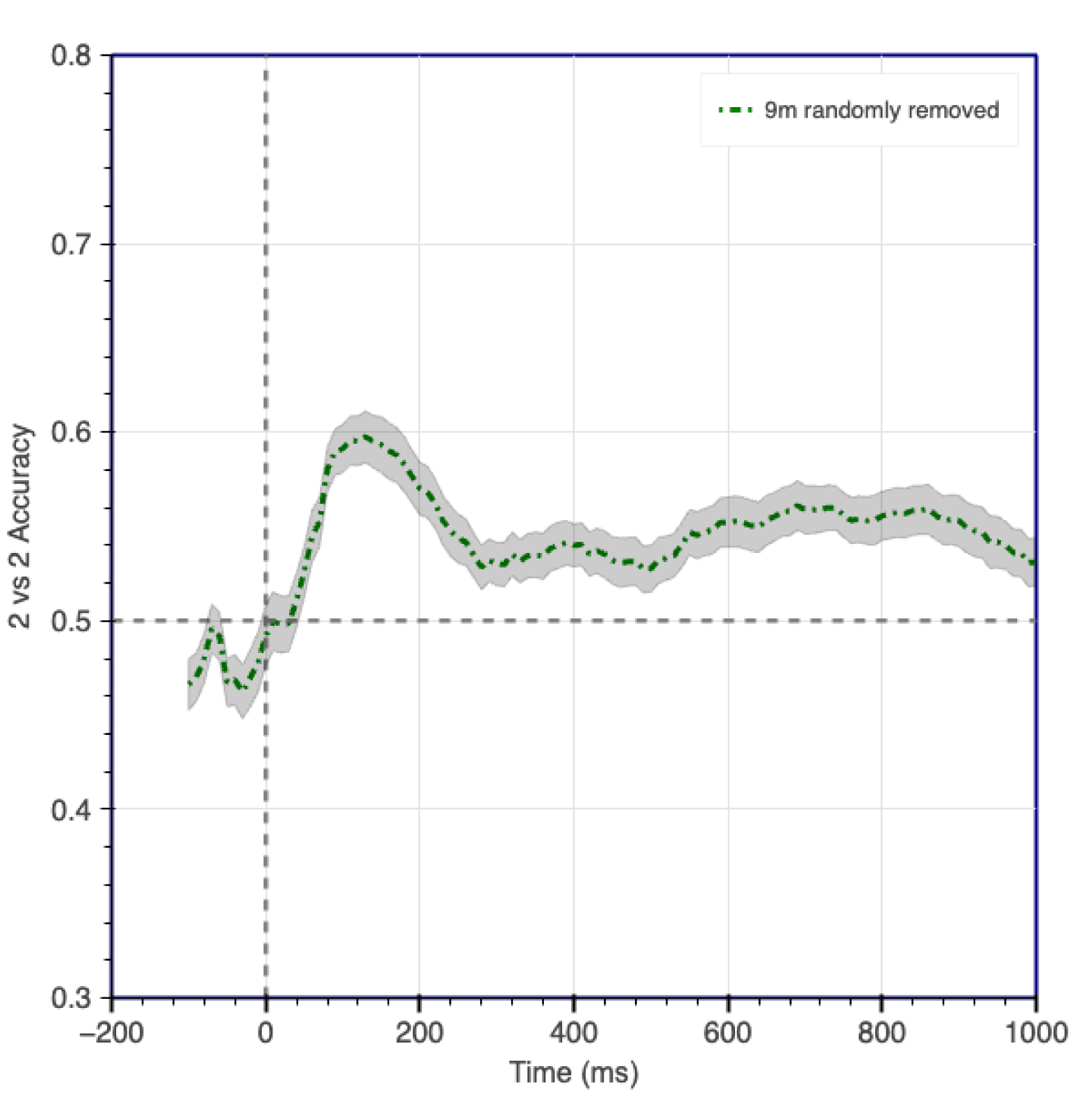
Word2Vec decoding results (Analysis 2) for the 9-month-old group, when trials were randomly removed to match the number in the 12-month-old group. We observe that the dynamics of the neural representations is unchanged compared to when all the trials were included (Analysis 2). This indicates that the effect observed is not attributable to the excess number of trials in the 9-month-old group.

## Footnotes

2 https://code.google.com/archive/p/word2vec/

3 https://scikit-learn.org/stable/modules/generated/sklearn.model_selection.GridSearchCV.html

4 To account for the difference in the number of trials between the two age groups, we randomly removed trials from the 9-month-old group and then conducted the decoding analysis. However, we found no significant difference in the results after removing the trials. Semantic decoding results for the 9-month-old are provided in Fig S3.

5 https://en.wikipedia.org/wiki/International_Phonetic_Alphabet

6 https://www.dictionary.com/

7 https://www.wiktionary.org/

8 In order to investigate whether the stimuli words can be distinguished simply based on the word phonemes, we trained a simple ridge regression model on the phonetic vectors to predict the semantic vectors obtained from Word2Vec. We found that the performance of the model was not better than chance. The results in Analysis 3 thus help us observe the decodability of word semantic representations in the absence of phonetic information. This effect is especially apparent in the 12-month-old group.

9 https://radimrehurek.com/gensim/models/word2vec.html#gensim.models.word2vec.Word2Vec.train

10 https://mne.tools/stable/generated/mne.viz.plot_topomap.html

